# Decision Confidence and Outcome Variability Optimally Regulate Separate Aspects of Hyperparameter Setting

**DOI:** 10.1101/2024.10.03.616475

**Authors:** Kobe Desender, Tom Verguts

## Abstract

Reinforcement learning models describe how agents learn about the world (value learning), and how they interact with their environment based on the learned information (decision policy). As in any optimization problem, it is important to set the process hyperparameters, a process which also is thought to be learned (meta-learning). Here, we test a key prediction of meta-learning frameworks, namely that there exist one or more meta-signals that govern hyperparameter setting. Specifically, we test whether decision confidence, in a context of varying outcome variability, informs hyperparameter setting. Participants performed a 2-armed bandit task with confidence ratings. Model comparison shows that confidence and outcome variability are differentially involved in hyperparameter setting. A high level of confidence in the previous choice decreased hyperparameter setting of decision noise on the current trial: when a trial was made with low confidence, the choice on the next trial tended to be more explorative (i.e. high decision noise). Outcome variability influenced another hyperparameter, the learning rate for positive prediction errors (thus affecting value learning). Both strategies are rational approaches that maximize earnings at different temporal loci: the modulation by confidence causes more frequent exploration early after a change point, the modulation by outcome variability is advantageous late after a change point. Finally, we show that (reported) confidence in value-based choices reflects the action value of the chosen option (irrespective of the unchosen value). In sum, decision confidence and outcome variability reflect distinct signals that optimally guide the setting of hyperparameters in decision policy and value learning, respectively.

## Introduction

In a complex and constantly changing environment, humans have to continuously update their beliefs about the environment and act upon these beliefs accordingly. Inflexibility to adapt and update beliefs leads to rigid behavior. For example, rather than always doing groceries at the same supermarket, it might be beneficial to explore other stores to see whether these have more qualitative or profitable offers. This is particularly the case in volatile environments, for example when a new owner is taking over. In such case, it might be optimal to explore and learn about alternative options. This example describes an intricate and delicate interplay between two key processes: learning about the value of stimuli in a changing environment (e.g. how good is this supermarket?) and acting based on these values (e.g. which supermarket will I frequent?). Reinforcement learning provides a mathematical framework to disentangle and study these two processes, by describing how humans estimate their beliefs (i.e. value learning) and subsequently make choices based on those beliefs (i.e. decision policy; Sutton & Barto, 2018). In the current work, we investigate whether, how and why feelings of decision confidence contribute to the optimization of value learning and decision policy.

### Value learning

Value learning is typically modeled as a process whereby action values are updated based on environmental feedback (Rescorla & Wagner, 1972; Sutton & Barto, 2018). These action values (often referred to as Q values) can be thought of as parameters that are updated in the face of new information. When an agent expects a large reward but receives nothing, a large (negative) prediction error follows, and the agent will update its action values based on this prediction error. The magnitude of this update is thought to be under control of a learning rate, which is a hyperparameter that determines how strongly action values are updated. If an agent sets a very high learning rate, action values will be updated very strongly in response to (unexpected) environmental feedback. Previous work has shown that in contexts of high versus low volatility, participants appropriately implement low versus high learning rates (Behrens et al., 2007). An active point of discussion is whether a single learning rate suffices for different type of prediction errors. For example, people tend to update their action values more strongly in response to positive prediction errors than in response to negative prediction errors, a finding that has been interpreted as a form of confirmation bias (Palminteri & Lebreton, 2022).

Remarkably, although having different learning rates for positive and negative prediction errors is sometimes described as a bias (Sharot et al., 2011), it might in fact be optimal to use higher learning rates for positive prediction errors (Lefebvre et al., 2022). In a similar vein, it has been debated whether humans merely update their beliefs about the value of the chosen option, or whether they also update their beliefs about unchosen options and if so whether this occurs via dissociable learning rates (Katahira, 2018; Palminteri et al., 2017). Combining both these features, this creates the theoretical possibility that humans actually use up to four different learning rates (obtained from crossing positive versus negative predictions error with updates about the chosen versus unchosen option).

### Decision Policy

Next to value learning, a process needs to be specified how to use such values to act, i.e. the decision policy. A straightforward policy is to plug the action values into a softmax decision rule, which then outputs a probability that an option will be selected (Daw et al., 2006). Such a decision policy is under the control of a hyperparameter, namely decision noise (i.e. inverse temperature) which controls the stochasticity of the choice. If an agent sets a very high level of decision noise, they will make very noisy decisions that are not so strongly driven by their action values. High decision noise effectively increases the probability of exploring options with lower values (Domenech et al., 2020; Jepma & Nieuwenhuis, 2011; Wilson et al., 2014). Instead, low decision noise causes exploitation because it effectively increases the probability of selecting the option with the highest value. Decision noise has been shown to depend on the choice horizon (Wilson et al., 2014), and baseline pupil diameter (Jepma & Nieuwenhuis, 2011).

### Meta-learning

Computational models based on the above two principles (value learning and decision policy) have proven to yield a highly successful algorithmic description accounting for human instrumental learning (Niv, 2009). However, although such models postulate *which* hyperparameters control human behavior, a key unresolved question is *how* and *why* an agent decides on the specific values of these hyperparameters. To address this question, meta-learning frameworks propose that the setting of hyperparameters is a process that itself is learned (Binz et al., 2023; Doya, 2002; Silvetti et al., 2018). Although learning rates are often modelled as stationary, there is evidence that these are not fixed but instead are dynamically adjusted depending on the volatility in the environment (Behrens et al., 2007). Learning to perform a task in a high versus low volatility context leads participants to use high versus low learning rates in a subsequent test phase (Simoens et al., 2024; Wen et al., 2023). In a similar vein, outcome variability also influences the learning rate (Diederen & Schultz, 2015; Preuschoff & Bossaerts, 2007). When outcomes are highly variable, agents (learn to) set a lower learning rate in order to avoid that action values are too heavily updated in response to noisy outcomes. Both findings (dependence on volatility and outcome variability) are consistent with the notion of a Kalman filter (Dayan et al., 2000). Here, the learning rate depends both on the level of outcome variability (i.e. expected uncertainty) and on the level of volatility (i.e. unexpected uncertainty; Soltani & Izquierdo, 2019). In a similar vein, although the decision policy is often modeled as a fixed parameter, recent evidence suggests instead that agents arbitrate between exploration and exploitation based on the confidence they have in their value representations (Boldt et al., 2019). Such a strategy is consistent with the idea of simulated annealing, which entails that an agent uses a high level of noise early in learning (when value representations are uncertain) and then slowly decreases the amount of noise as learning evolves (when value representations are more stable; Cagan & Kotovsky, 1997).

### Which signal(s) drive(s) meta-learning?

Thus, reinforcement learning parameters may be meta-learned as a function of the environment the agent is in. It follows that the brain needs “meta-signals” to set these hyperparameters. Similar to how the brain uses prediction errors (i.e. between expected and obtained rewards) as a signal to update its action values, it also requires some meta-signal to update its hyperparameters. At current, however, there is no clear consensus regarding the nature of these meta-signals. As described above, learning rates may depend on outcome variability and environmental volatility, but it is unclear whether and how agents represent these different forms of variability. One prime candidate meta-signal is the sense of confidence that typically accompanies a decision. Indeed, confidence has been shown to act as a metacognitive regulatory signal affecting learning (Cortese, 2022; Drugowitsch et al., 2019; Lak et al., 2020; Meyniel & Dehaene, 2016), task prioritization (Aguilar-Lleyda et al., 2020), information seeking (Desender et al., 2018; Schulz et al., 2023), and comparison of performance between different tasks (de Gardelle & Mamassian, 2014). Moreover, (outcome) variability is known to influence subjective confidence (Boldt et al., 2017; Spence et al., 2016). Whereas outcome variability describes an experimental manipulation from the experimenter’s point of view, subjective confidence about value-based decisions reflects a subjective perception from the participants’ point of view. Therefore, it is theoretically possible that outcome variability steers the hyperparameters controlling learning only via its effect on confidence. However, other work has provided suggestive evidence that confidence in value beliefs might balance the tradeoff between exploration and exploitation (Boldt et al., 2019). The conceptual definition of confidence in two-armed bandit tasks has received relatively little attention compared to the perceptual domain (for an exception, see Salem-Garcia et al., 2023a). Whereas a growing consensus is that confidence in perceptual decisions can be modelled as the probability of a choice being correct (Calder-Travis et al., 2024; Fleming & Daw, 2016; Kiani et al., 2014; Le Denmat et al., 2024; Sanders et al., 2016), it is less clear whether the same frameworks can be used to understand value-based confidence. In sum, it remains unclear what confidence reflects in the context of value-based learning, and whether it serves the role of a meta-signal controlling hyperparameters setting.

### The current work

To unravel whether decision confidence acts as a meta-signal informing the process of hyperparameter setting during meta-learning and how both relate to outcome variability, we directly manipulated outcome variability and measured its effect on confidence in a two-armed bandit task. First, we tested whether confidence in value-based decisions can be understood as the probability of being correct (i.e. selecting the high-value bandit) conditional on the available evidence. Using computational modeling, we then examined the influence of confidence and outcome variability on value learning and decision policy. Outcome variability was manipulated by sampling rewards from a very narrow or a very wide distribution. The association between slot machine and reward magnitude changed unexpectedly during the course of the experiment. Decision confidence was expected to depend on this manipulation of (reward) variability, as well as to be sensitive to these sudden reversals. Using this design, the current study allows to assess whether confidence reflects a meta-signal that is used for hyperparameter setting. In order to do so, we developed a regression approach where, instead of fixing hyperparameters to a single value, these were allowed to dynamically co-vary with fluctuations in the predictor variables (i.e. outcome variability and confidence). Finally, to assess whether we could see durable changes in parameter setting, we induced different levels of outcome variability during an induction phase in which participants could meta-learn the optimal strategy, and we then tested the transfer of this strategy in an unbiased testing phase in which outcome variability was matched (i.e. a transfer effect). Using this approach, the current study addresses whether and how either outcome variability and confidence interact with setting value learning and decision policy meta-parameters.

## Methods

### Participants

The experiment comprised sixty participants, all of which were first year psychology students at the KU Leuven who participated in return for course credits. Data of six participants was removed because performance was at chance level in at least one of the blocks, thus the final sample comprised fifty-four participants. Each participant took part in two one-hour sessions on separate days, separated at least 12h and less than 72h. Participants carried out the task on their own pc, with the data being collected online via the pavlovia platform (pavlovia.org). Only participants with a Windows PC with an external mouse were allowed to participate. The three participants with the highest score received a gift voucher from a local store. Information about age, gender and handedness was not saved and is therefore unknown, but in this participant pool a majority of right-handed, 18-year old mostly females can be expected. All participants provided an informed consent via e-mail and were naive regarding the hypotheses of the study. The study was approved by the Social and Societal Ethics committee of the KU Leuven (SMEC).

### Materials and procedure

The experiment was programmed in Psychopy Builder (Pierce et al., 2019). Participants performed a two-armed bandit task with probability distributions of the rewards changing over time. On each trial, participants decided between two slot machines that were always presented left and right on the screen by pressing the “s” or “f” on the keyboard, respectively. After their choice, a horizontal confidence scale was presented on the screen with the labels “this was a guess” to “very certain” at the left and right outer ends, respectively. Participants indicated their level of confidence in the choice they made by clicking with the mouse on the corresponding location of the continuous confidence scale. Confidence ratings were scored between 1 (“this was a guess”) and 5 (“very certain”). Finally, a reward was shown centrally on the screen in the form of a number presented inside a star (see Figure 1A).

**Figure 1.**
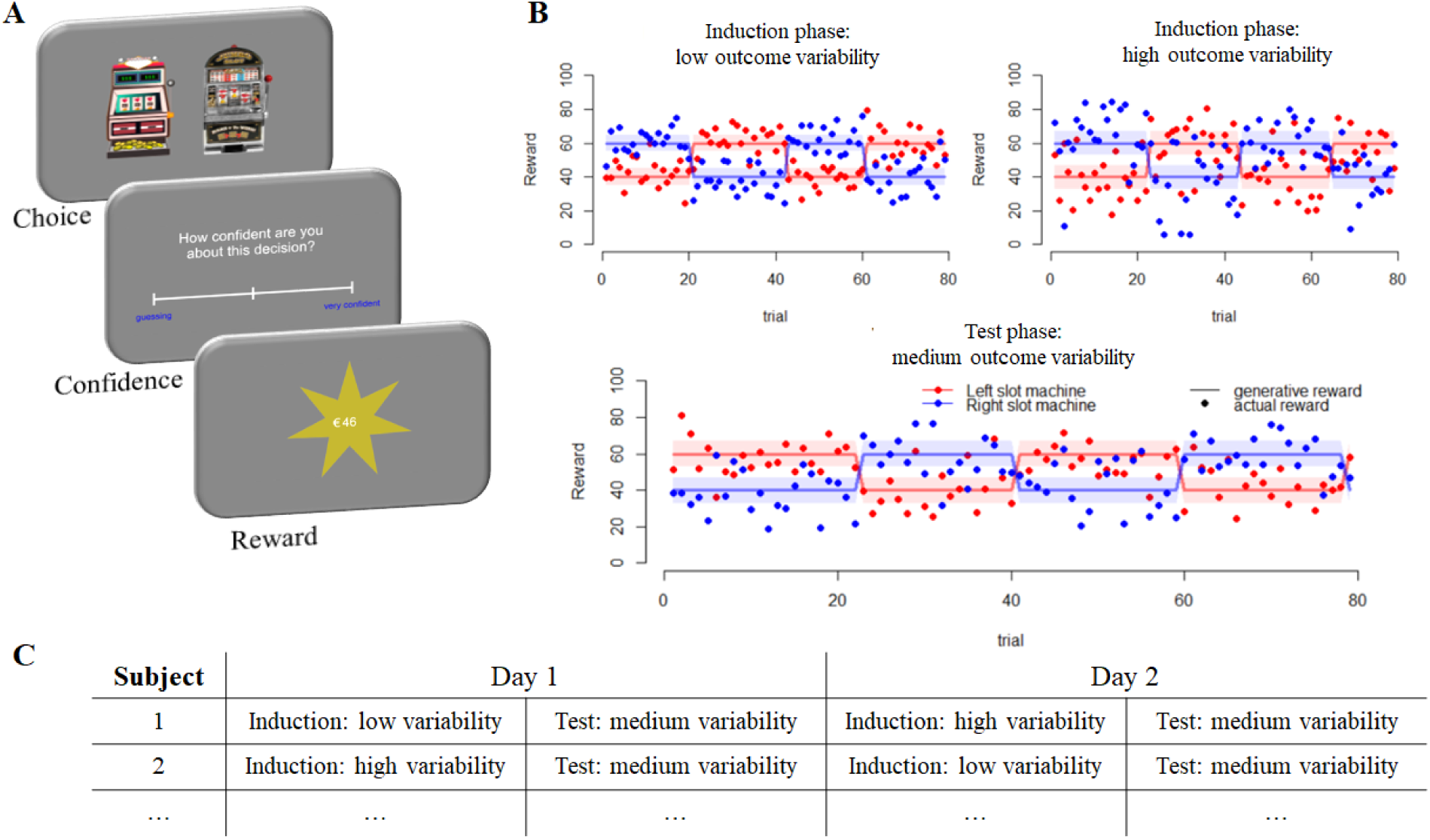
Experimental Paradigm. **A.** On each trial participants decided between two slot machines by pressing “s” or “f” and then indicated their level of confidence in this choice using the mouse on a continuous slider running from “This was a guess” to “Very Certain”, which was coded between 1 and 5, respectively. Afterwards they were shown how much reward they obtained on that trial. **B.** Rewards were drawn from a high (M=60) or low (M=40) generative mean, with switches every 18-22 trials regarding the association between slot machines and generative means. During an induction phase, outcome variability was manipulated by drawing rewards from a low-variable distribution (SD=8, left panel) or a high-variable distribution (SD=16, right panel). During a test phase rewards were drawn from an intermediate distribution (medium-variability, SD=12, bottom panel). Dots show illustrative examples, lines show the generative mean and shades show the generative variability. **C.** The experimental design was fully within-subjects. On separate days, participants performed the induction phase with high or low outcome variability (order counterbalanced between participants), followed by the test phase with medium outcome variability.

The reward depended on the slot machine chosen. Rewards were sampled from a distribution with a high mean (*M*=60) for one slot machine and a low mean (*M*=40) for the other slot machine. The association between slot machines and reward mean switched every 18-22 trials (see Figure 1B). Participants were informed that the association between slot machine and reward would frequently switch. We manipulated outcome variability in two separate sessions. In one of the sessions reward outcomes were sampled from a low variance distribution (SD=8), whereas in the other session reward outcomes were sampled from a high variance distribution (SD=16). In the test phase of both sessions reward outcome was sampled from a medium-variance distribution (SD=12).

The experimental manipulations were fully within-subjects. Half of the participants started session 1 with 400 trials of the low outcome variability condition (induction phase), followed by 400 trials of the medium outcome variability condition (test phase); and started session 2 with 400 trials of the high outcome variability condition (induction phase), followed by 400 trials of the medium outcome variability condition (test phase). For the other half of the participants everything was the same except that they started with the high outcome variability condition during session 1 (see Figure 1C).

### Statistical analyses

Behavioral data were analyzed using linear mixed effects models using the lme4 package (Bates et al., 2015). We fitted random intercepts for each participant; error variance caused by between-subject differences was accounted for by adding random slopes to the model. The latter was done only when this significantly increased the model fit, as assessed by model comparison. Confidence reports were analyzed using linear mixed models, for which F statistics are reported and the degrees of freedom were estimated by Satterthwaite’s approximation (Kuznetsova et al., 2014). Choice data was analyzed using logistic linear mixed models, for which χ2 statistics are reported. Significant interactions were followed-up by computing contrasts using the glht package. For non-significant effects of importance we additionally computed a Bayes Factor using BayesFactor package using the default priors (Morey & Rouder, 2014). Model fitting was done in R (R-Core-Team, 2016). Mediation analysis was performed using the mediation package (Tingley et al., 2014). A mediator mixed model was fitted predicting confidence based on outcome variability with random slopes for outcome variability, and an outcome model was fitted predicting next-trial choice accuracy based on confidence and outcome variability with random slopes for confidence. A mediation analysis was then performed with these two models, testing whether the influence of outcome variability on next-trial choice accuracy was mediated by confidence.

### Reinforcement learning model

To explain behavior in the two-armed bandit task we assumed that participants learned the action value, *Q*, of option *k* at time 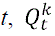, for each slot machine by updating it on each trial in response to reward, *R_t_*, using the following updating rule:

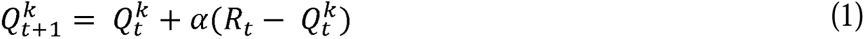

with *α* reflecting the learning rate. To make a choice between the two slot machines, we assume that participants select slot machine *k* with probability:

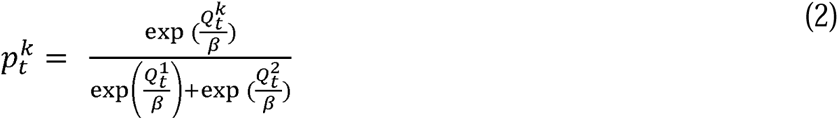

with *β* reflecting a decision noise parameter controlling the level of stochasticity in the choice (also referred to as the temperature parameter). Model fitting was done separately for each participant, in R using the DEoptim library (Mullen et al., 2011), by minimizing the negative log likelihood. Our modeling approach consisted of two separate steps. In a first step, we compared the fit of different models of increasing complexity (while ignoring potential effects of outcome variability and confidence). We started with the basic RL model described above, featuring one level of decision noise *β* and a single learning rate *α* that was used to update the value of the chosen option only (Model 0, two free parameters). This model was then compared to Model 1 in which both chosen and unchosen value were updated using the same prediction error (but in different directions; Model 1, two free parameters), a model with separate learning rates for the chosen and unchosen value (Model 2, three free parameters), a model with separate learning rates for positive and negative prediction errors (Model 3, three free parameters), and a model with separate learning rates for each combination of chosen vs unchosen and positive vs negative prediction errors (Model 4, five free parameters). See Table 1 for an overview of the different models and their fit to the data.

**Table 1.** Overview of the different models and their associated fit. During the first step, models with varying complexity were tested in their ability to fit the data. In a second step, we assessed the effect of outcome variability by testing for each parameter in Model 5 whether the fit improved if this parameter was separately estimated for low and high outcome variability, indicated by the subscripts low and high, respectively. To assess the effect of decision confidence, we tested for each parameter in Model 5 whether the fit improved if the hyperparameter was estimated using an intercept w0 and a slope w1 relating previous-trial confidence to that hyperparameter. Winning models per section (i.e. lowest BIC) are indicated in bold. Note, α and β refer to learning rate and decision noise, and subscripts refer to chosen option, C, unchosen option, U, positive prediction error, +, and negative prediction error, −.

| Identifying the best fitting model |  |  |  |  |  |  |  |  |  |  |  |  |
| --- | --- | --- | --- | --- | --- | --- | --- | --- | --- | --- | --- | --- |
| Model | $\alpha$ | $\alpha_C$ | $\alpha_U$ | $\alpha_+$ | $\alpha_-$ | $\alpha_{C+}$ | $\alpha_{C-}$ | $\alpha_{U+}$ | $\alpha_{U-}$ | $\beta$ | df | BIC |
| 0 | - | ✓ | - | - | - | - | - | - | - | ✓ | 2 | 1045 |
| 1 | ✓ | - | - | - | - | - | - | - | - | ✓ | 2 | 980 |
| 2 | - | ✓ | ✓ | - | - | - | - | - | - | ✓ | 3 | 911 |
| 3 | - | - | - | ✓ | ✓ | - | - | - | - | ✓ | 3 | 969 |
| 4 | - | - | - | - | - | ✓ | ✓ | ✓ | ✓ | ✓ | 5 | 816 |
| 5 | - | - | ✓ | - | - | ✓ | ✓ | - | - | ✓ | 4 | **817** |
| The effect of outcome variability |  |  |  |  |  |  |  |  |  |  |  |  |
| Model | $\alpha_{C+}$ | | $\alpha_{C-}$ | | $\alpha_U$ | | $\beta$ | | | | | |
| 6 | ✓<sub>low</sub> | ✓<sub>high</sub> | ✓ |  | ✓ |  | ✓ |  |  |  | 5 | **812** |
| 7 |  | ✓ | ✓<sub>low</sub> | ✓<sub>high</sub> | ✓ |  | ✓ |  |  |  | 5 | 816 |
| 8 |  | ✓ |  | ✓ | ✓<sub>low</sub> | ✓<sub>high</sub> | ✓ |  |  |  | 5 | 817 |
| 9 |  | ✓ |  | ✓ |  | ✓ | ✓<sub>low</sub> | ✓<sub>high</sub> |  |  | 5 | 824 |
| The effect of decision confidence |  |  |  |  |  |  |  |  |  |  |  |  |
| Model | $\alpha_{C+}$ | | $\alpha_{C-}$ | | $\alpha_U$ | | $\beta$ | | | | | |
| 10 | ✓<sub>w0</sub> | ✓<sub>w1</sub> | ✓ |  | ✓ |  | ✓ |  |  |  | 5 | 815 |
| 11 |  | ✓ | ✓<sub>w0</sub> | ✓<sub>w1</sub> | ✓ |  | ✓ |  |  |  | 5 | 811 |
| 12 |  | ✓ |  | ✓ | ✓<sub>w0</sub> | ✓<sub>w1</sub> | ✓ |  |  |  | 5 | 818 |
| 13 |  | ✓ |  | ✓ |  | ✓ | ✓<sub>w0</sub> | ✓<sub>w1</sub> |  |  | 5 | **805** |
| Combining outcome variability and confidence |  |  |  |  |  |  |  |  |  |  |  |  |
| Model | $\alpha_{C+}$ | | $\alpha_{C-}$ | | $\alpha_U$ | | $\beta$ | | | | | |
| 14 | ✓<sub>low</sub> | ✓<sub>high</sub> | ✓ |  | ✓ |  | ✓<sub>w0</sub> | ✓<sub>w1</sub> |  |  | 6 | **799** |

After identifying the best fitting model, in a second step we investigated the role of outcome variability and confidence on each of the parameters. Specifically, we selected the winning model of step 1, and then tested for each of the parameters in the winning model whether it was modulated by outcome variability (by estimating the parameter separately for high vs low outcome variability conditions). Next, we also tested for each of the parameters in the winning model whether it was modulated by the reported level of decision confidence. Because confidence varied continuously, the latter was done by testing for each parameter whether it depends on previous-trial confidence. Specifically, instead of a single constant value for a parameter (say *β*), instead we tested whether model fit improved when the parameter was quantified at trial *t* using the following equation:

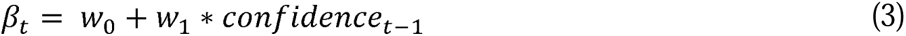

If the estimate of *w*_1_ is significantly different from zero, this would be evidence that previous-trial confidence influences the current level of decision noise. Note that the example here describes the approach for *β* only, but this was implemented for all of the parameters in the winning model of step 1.

### Modeling confidence in an RL model

Up until this point, we only modeled choices using our computational model, whereas confidence simply was the value reported by participants. To explain how reported confidence arises during value-based learning, we next adapted our model to account for confidence judgments. We compared three nested models in their ability to explain empirical confidence judgments: a model in which confidence is based on Q_chosen,t_, a model in which confidence is based on Q_chosen,t_-Q_unchosen,t_, and a model in which confidence is based on (Q_chosen,t_-Q_unchosen,t_) *β*_1_ (note, these values were taken from Model 14 so *β* was not constant but varied on a single-trial basis). Importantly, the previous section showed an influence of *previous-trial* confidence on current trial decision policy, whereas here we model the *current-trial* confidence as a function of the current-trial Q values. Given this temporal difference there is no circularity in our analysis. For each of the three models, we took the estimates of the best fitting model from the previous stage (i.e., Model 14; i.e. parameters were not fitted again). In order to map the model-based quantity onto the five-point scale used by participants, we additionally estimated four confidence thresholds. Note that the three models have the same number of free parameters and so we directly compared their residual error, RSS, in order to compare model fit. The four parameters were optimized to minimize the following cost function:

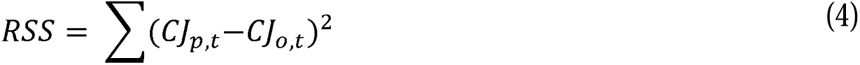

where CJ reflects average confidence judgment at trial *t,* reflecting the first 15 trials after a switch, and *p* and *o* referring to predicted and observed data, respectively.

## Results

### The effect of outcome variability on accuracy and confidence

To examine the influence of outcome variability on accuracy, we used a logistic mixed model to test whether the probability of selecting the correct slot machine depends on outcome variability (2 levels: high vs low outcome variability), experiment phase (2 levels: induction vs test), and their interaction. The model included a full random effects structure. Note that outcome variability was identical in the two test phases (i.e. at medium level) such that the simple main effect of outcome variability in the test phase reflects a carry-over effect from the induction phase (i.e. meta-learning). The main effect of outcome variability, χ2(1) = 32.59, *p* < .001, showed that people more frequently selected the high-reward bandit when outcome variability was low (*M* = 84.8%) compared to when it was high (*M* = 79.1%). The data also showed a main effect of phase (i.e. a slight decrease in selecting the high-reward bandit from induction to test), χ2(1) = 20.32, *p* < .001, as well as an interaction between both, χ2(1) = 23.89, *p* < .001. Critically the effect of outcome variability was significant both in the induction phase (87.1% vs 78.6%), *z* = 7.22, *p* < .001, as well as in the test phase (82.2% vs 79.6%), *z* = 3.42, *p* < .001. Given that outcome variability was at medium level during the test phase, this finding suggests that participants meta-learned a strategy during the induction phase that they continued to use to some extent during the (unbiased) test phase (Figure 2A-B).

**Figure 2.**
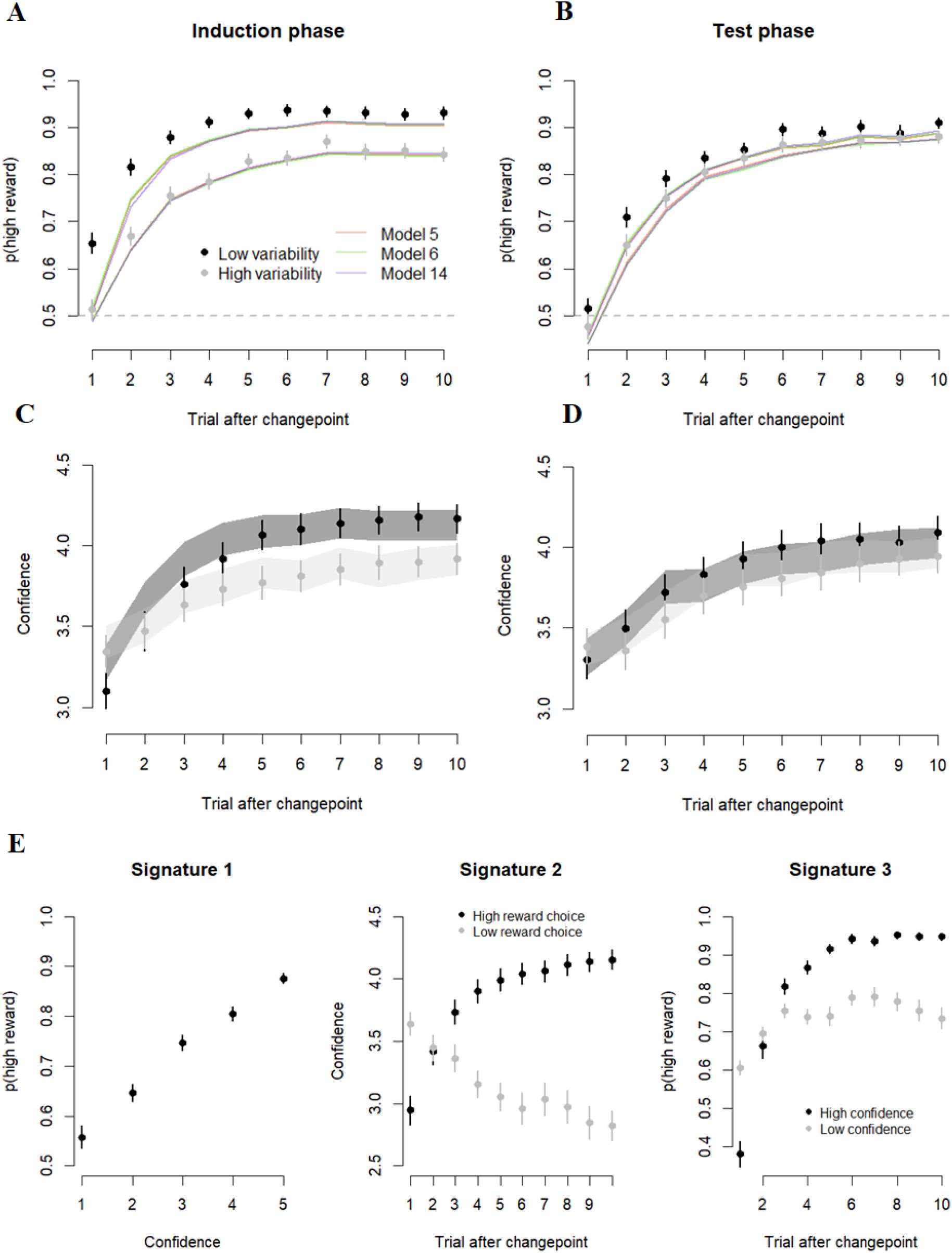
Behavioral results of the two-armed bandit task. A-B. Learning curves. After a change point, participants rapidly adapted to the change in contingencies and quickly started to prefer the high reward slot machine. The learning curves were steeper when outcome variability was low versus high, both during the induction phase (A) and during the test phase in which outcome variability was the same for both (B). **C-D. Confidence learning curves.** The reported level of decision confidence closely mirrored the learning curves, with a rapid increase in confidence right after a change point. As expected, confidence was higher when outcome variability was low versus high. **E. Confidence signatures.** Confidence in value-based decisions displays the three qualitative signatures that are characteristic of confidence in perceptual and memory-based decision making. Note: points and error bars show the mean and SEM of the empirical data; shaded lines show the model fit, with the width reflecting SEM. Note that in A-B SEM of model fit is not shown to avoid clutter.

Results for confidence closely mirrored the choice findings (Figure 2C-D). The main effect of outcome variability, χ2(1) = 5.44, *p* = .019, showed that confidence was higher when outcome variability was low (*M* = 3.94) compared to when it was high (*M* = 3.78). Although the main effect of phase was not significant, *p* = .227, there was a significant interaction, χ2(1) = 28.08, *p* < .001. Outcome variability influenced confidence both in the induction phase (3.98 vs 3.79), *z* = 2.89, *p* = .004, as well as numerically in the test phase (3.89 vs 3.78), *z* = 1.75, *p* = .081. Note that this analysis underestimates the influence of outcome variability on confidence, because it does not consider how far apart trials are from a change point, whereas in Figure 2B and 2D it can be seen that the influence of outcome variability on confidence is rather weak (or even reversed) in the first couple of trials following a change point, whereas later on the effect is more consistent.

### Confidence in value-based decisions tracks the probability of being correct

The observation that the learning curves for confidence closely mirror the learning curves for accuracy already suggests that confidence in value-based decisions reflects the probability of being correct (i.e. selecting the high-value bandit), similar to perceptual decisions. To more formally corroborate this notion, we computed three well-known signatures of statistical decision confidence in perceptual and memory decisions (Sanders et al., 2016). The first signature reflects a monotonic relation between confidence and accuracy (i.e. in this case selecting the high-reward option). Figure 2E shows that confidence (binned in five equal-sized bins) monotonically relates to the proportion of high-reward choices, χ2(1) = 229.27, *p* < .001, with significant differences between all adjacent bins, *z*s > 9.06, *p*s < .001. The second signature reflects a positive relation between evidence and confidence for correct trials, but a negative relation for error trials. Although the concept of evidence in perceptual decision making does not have a one-to-one mapping with evidence in value-based decisions, we here assumed that evidence increases with trial number (which is in line with the finding that accuracy increases with trial number), and so we take trial number as a proxy for evidence. When doing so, we see the typical “folded-X pattern” (see Figure 2E). A mixed model predicting confidence based on trial, high reward choice and their interaction indeed showed a significant interaction, χ2(1) = 3164.85, *p* < .001, reflecting that confidence increased with trial number for high-reward choices, *b* = .07, *z* = 56.26, *p* < .001, and decreased for low-reward choices, *b* = –.04, *z* = –40.35, *p* < .001. Finally, the third signature reflects a steeper psychometric curve (high-reward choice as a function of trial number) for high versus low confidence trials. We thus tested whether the increase in high-reward choices across trials was steeper for high than for low confidence trials. Indeed, a logistic mixed model predicting high-reward choices by trial number, a binary measure of confidence (2 levels: low vs high) and their interaction showed a significant interaction effect, χ2(1) = 1410.05, *p* < .001, reflecting a stronger association between trial number and high-reward choices for trials judged with high confidence, *b* = .15, *z* = 38.17, *p* < .001, compared to trials with low confidence, *b* = .06, *z* = 5.39, *p* < .001. We conclude that decision confidence in value-based decision making behaves highly similar to confidence in perceptual decision-making tasks, confirming that decision confidence in such tasks reflects the probability of having selected the high reward option.

### Building a model to explain choices

To explain the computations underlying the choices that participants made, we resorted to RL principles. In a first step, we constructed a model that provided the best fit to the data. These models were fitted on all data from both the induction and the test phase. Results from model fits performed separately for each phase are reported below in a designated section. Details of all models as well as their corresponding fit can be found in Table 1. We started with a standard RL model (Model 0) in which the value for the chosen (but not the unchosen) slot machine is updated based on the prediction error scaled by learning rate α, and a noise parameter β controls the policy decision noise (two free parameters). Next, given that participants were informed at the start of the experiment that the two bandits were anti-correlated, we considered it reasonable that they used the prediction error to not only update the value of the chosen option, but also the value of the unchosen option. Indeed, Model 1, in which both chosen and unchosen value were updated using the same prediction error (but in different directions) scaled by the same learning rate provided a better fit than Model 0 in terms of BIC (*M* = 1045 vs *M* = 980). We then compared performance of Model 1 to models with separate learning rates for the chosen and unchosen value (Model 2; three free parameters); a model with separate learning rates for positive and negative prediction errors (Model 3; three free parameters); and separate learning rates for each combination of chosen vs unchosen and positive vs negative prediction errors (Model 4; five free parameters). Model 4 provided the best fit to the data (ΔBICs > 95; Figure 3A). Closer inspection of this model revealed a significant difference between learning rates for positive versus negative prediction errors for the chosen value, *t*(53) = 2.92, *p* = .005, but not for the unchosen value, *p* = .149.

**Figure 3.**
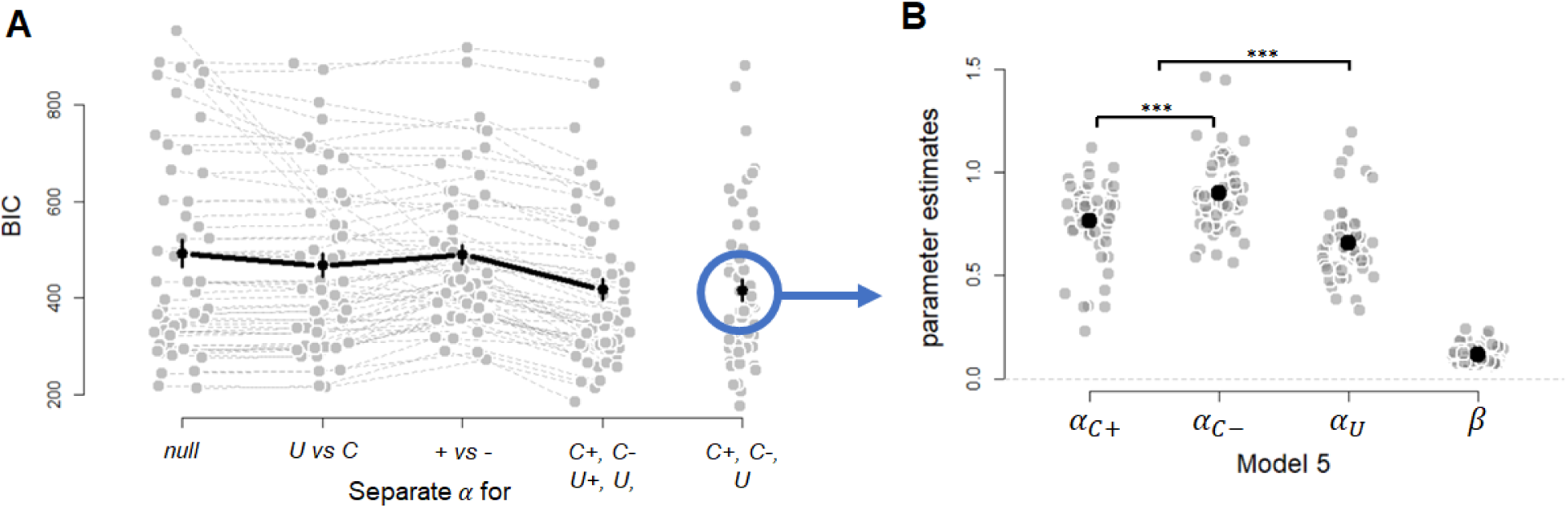
Results of the first step, identifying the best fitting model. **A**. We compared the fit of model variants that differed in whether they had separate learning rates for positive (+) and negative (−) prediction errors and the chosen (C) and unchosen (U) option. **B.** The winning model featured separate learning rates for positive and negative prediction errors for the chosen option, α_c+_ and α_c−_respectively, a single learning rate for the unchosen option, α_u_, and α decision noise parameter, β. Note, subscripts refer to chosen option (C), unchosen option (U), positive prediction error (+), and negative prediction error (−). Grey dots show individual participants, same conventions as in Figure 2.

Thus, for parsimony we additionally fitted Model 5, in which separate learning rates were estimated for positive versus negative predictions errors for the chosen value, but a shared (single) learning rate for the unchosen value (irrespective of prediction error sign). As expected, this model showed a significant difference between learning rates for positive versus negative prediction errors for the chosen value, *t*(53) = 3.82, *p* < .001, and it also showed that the learning rate for the unchosen option was lower than the learning rate of the chosen options, *t*(53) = 5.60, *p* < .001 (see Figure 3B). Importantly, Model 5 provided a similar fit to the data as Model 4 (ΔBIC = –1), effectively gaining a free parameter without sacrificing model fit. The fits of Model 5 are visualized in Figure 2A and 2B. Note that while the fit for the high variability condition provided a very close match to the data, the model slightly underestimates the learning curve in the low variability condition, potentially because some participants figured out that change points occur every ∼20 trials and started to anticipate these change points. For convenience, in the remainder of the paper we will omit ‘for the chosen value’ when describing the first two learning rates, and thus simply refer to “learning rates for positive and negative prediction errors”.

### Modeling the effect of outcome variability on value-based learning

Having constructed a good model to explain the choice data, we went on to the second step: investigating the role of our manipulation of outcome variability (high vs low) on value learning. To do so, we further built on Model 5, identified as the most parsimonious model to explain the performance data. Specifically, we estimated four additional models (Models 6 to 9, one for each parameter in Model 5) in which we tested whether model fit improved if we estimated that parameter separately for high and low outcome variability conditions (again collapsing data across induction and test phase). Relative to Model 5, a model with learning rates for positive prediction errors separately for the high and low outcome variability condition (Model 6) provided the best fit (ΔBIC = 5.00; see Figure 4A). In line with optimality considerations (Dayan et al., 2000), learning rates for positive prediction errors were larger in the low variability context (*M* = .79) than in the high variability context (*M* = .71), *t*(53) = 2.57, *p* = .013 (see Figure 4C). In contrast, estimating separate high and low variability estimates was not advantageous for the learning rate for negative prediction errors (ΔBIC = 1.46), for the learning rate for the unchosen value (ΔBIC = –0.22), and perhaps surprisingly also not for decision noise (ΔBIC = –7.03) (see Figure 4A).

**Figure 4.**
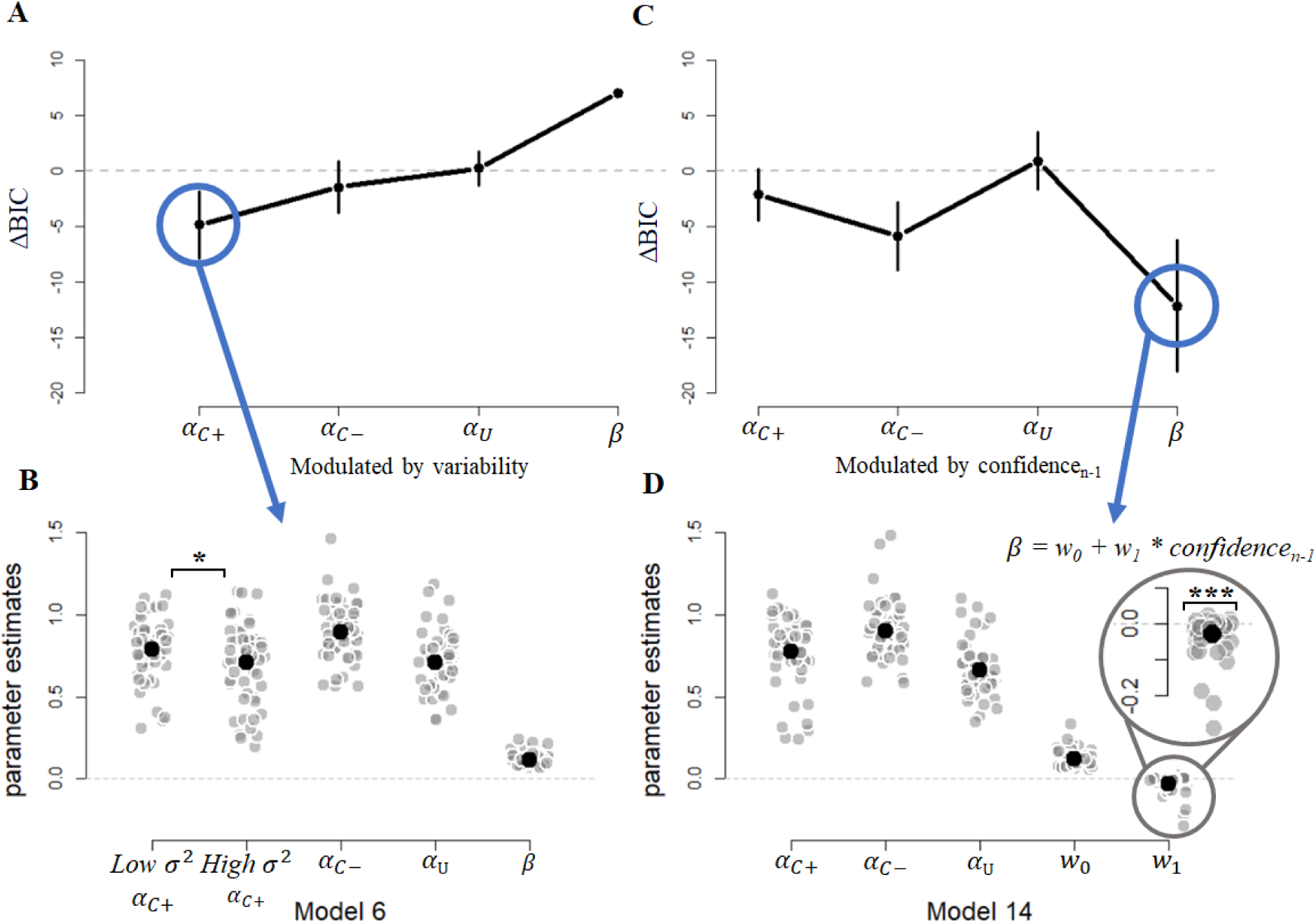
The influence of outcome variability and confidence on learning. **A.** For each parameter in Model 5, identified as the most parsimonious model during the model-building phase, we constructed additional models to test whether the fit improved if a parameter was estimated separately for high and low outcome variability (Models 6-9). The best model (Model 6) featured learning rates for positive prediction errors separately for high and low outcome variability. **B.** Parameter estimates of Model 6 confirmed that learning rates for positive prediction errors were significantly different when estimated separately for high and low outcome variability. **C.** We then applied the same strategy for confidence and tested whether model fit improved if the parameter was allowed to vary on a trial-by-trial basis as a function of previous-trial confidence. In the winning model (Model 14), decision noise was dependent on previous-trial confidence. **D.** Estimates from Model 14 confirm that the parameter controlling for the influence of previous-trial confidence on decision noise (w_1_) was significantly different from zero. Note: same conventions as in Figure 3.

### Consequences of confidence in value-based learning

After demonstrating that our data are best accounted for by a model in which the learning rate for positive prediction errors depends on outcome variability, we next repeated the same modeling procedure but now testing whether each of the parameters in the winning model were modulated by previous-trial confidence. To achieve this, we implemented a regression approach whereby model parameters were allowed to vary dynamically as a function of previous-trial confidence (in line with previous work relating previous-trial confidence to current trial DDM parameters; Desender et al., 2019; van den Berg et al., 2016). Given that outcome variability influences decision confidence (see the section reporting Choice and Confidence Behavior), one hypothesis is that decision confidence has the same effect as outcome variability: i.e. modulating the learning rate for positive prediction errors. However, as outlined in the introduction an alternative hypothesis is that confidence instead modulates the decision policy. To arbitrate between these hypotheses, for each of the parameters in Model 5, we tested whether model fit improved if the parameter was allowed to vary as a function of previous-trial decision confidence. Rather than estimating a single value of that parameter that remains constant throughout the experiment, instead we estimated an overall intercept, *w*_0_ (i.e. constant during the experiment), and a slope, *w*_1_, determining the extent to which previous-trial confidence modulates the parameter on a trial-by-trial basis (see Equation 4).

Interestingly, Model 10 in which the learning rate for positive prediction errors depends on previous-trial confidence did not show a clear improvement relative to Model 5 (ΔBIC = 2.14). Instead, the best model was Model 13, in which decision noise was allowed to vary as a function of previous-trial confidence (ΔBIC = 12.00; Figure 4B). Within this model, the estimate controlling the influence of previous-trial confidence on current-trial decision noise was significantly different from zero, *w*_1_ = –.029, *t*(53) = –3.78, *p* < .001 (Figure 4D). This negative slope reflects that higher values of previous-trial confidence are associated with lower decision noise. Finally, the increase in model fit was smaller when previous-trial confidence affects the learning rate of negative prediction errors (ΔBIC = 5.86) or unchosen options (ΔBIC = –0.88).

On every trial, participants were asked about their level of confidence in their choice before receiving a reward. As previously shown in Figure 2E, confidence closely corresponds to the probability of selecting the high-reward option (i.e. confidence signature 1), and hence there is a robust correlation between confidence and reward (average *r* = .12, *p* < .001). An alternative hypothesis is therefore that current trial decision noise is modulated by the feedback that participants saw, rather than their level of confidence. However, this alternative hypothesis is not supported by the data: a model in which previous-trial feedback influences decision noise provides a worse fit than Model 13 (ΔBIC = –15.00).

Finally, we fitted Model 14 in which we combined the effect of outcome variability on the learning rate for positive prediction errors, and the effect of confidence on decision noise in the same model. This model had the best fit of all models under consideration so far, being superior to both Model 6 (ΔBIC = 13) and Model 13 (ΔBIC = 6). As expected, in this model there was a significant difference between learning rates for positive prediction errors depending on outcome variability, *t*(53) = 2.69, *p* = .009, and a significant trial-by-trial relation between confidence and decision noise, *t*(53) = –3.78, *p* < .001.

It might seem surprising that although at the behavioral level we observed an influence of outcome variability on confidence, outcome variability and confidence have distinct effects on the parameters. This is particularly striking, given that in perceptual decision making it has been shown that the influence of variability on information seeking is mediated by confidence (Desender et al., 2018). Given this apparent contradiction, we ran a trial-based mediation analysis testing whether the influence of outcome variability (high or low) on choice behavior (selecting the high or low reward option) is mediated by confidence. We first confirmed that in the mediator model confidence was significantly predicted by outcome variability, *b* = .15, *t* = 2.34, *p* = .023, and in the outcome model next-trial choices were significantly predicted by outcome variability, *b* = .32, *z* = 18.05, *p* < .001, and by confidence, *b* = .21, *z* = 7.42, *p* < .001. A mediation analysis based on these models showed that 90.7% of the influence of outcome variability on choice went via the direct path (*p* < .001); whereas only 9.3% of this effect was mediated via confidence (*p* =.005). This analysis is in line with the notion that in value-based decision-making contexts outcome variability and confidence have largely independent effects on choice behavior.

### Modulating decision noise by previous-trial confidence and learning rate by outcome variability is the optimal strategy

In the previous section we demonstrated that participants rely on an internal sense of (previous-trial) confidence to determine their decision noise. Specifically, with low confidence they tend to make noisy decisions; with high confidence they tend to make less noisy decisions. Intuitively, this seems a sensible strategy because under low confidence it might be beneficial to explore other options (see Lefebvre et al., 2022 for a similar argument in the context of confirmation bias). In a similar vein, outcome variability modulated the learning rate for positive prediction errors, which likewise seems a sensible strategy because it avoids too strong updates when outcomes are noisy. To confirm both these intuitions, we simulated data separately for each participant based on the fitted parameters of the overall winning model (i.e. Model 14), which included both of these effects. Critically, during the simulations we systematically varied *w*_1_ between –.35 and .05 (steps of .0025), and we systematically varied *high σ^2^ α_c+_* – low *σ^2^ α_c+_* such that there was a difference of –.2 up to a .1 (steps of .01). For all these combinations, we computed the expected probability of selecting the high reward choice that would have been obtained if that level of *w*_1_ and *α_c+_* would have been used. Figure 5A shows this optimality surface, and as can be seen the point of optimality was negative for both, meaning that the optimal strategy was to be more explorative with low previous-trial confidence and to use a higher learning rate with low outcome variability. Having identified the optimal theoretical strategy, we next compared this to the empirical data.

**Figure 5.**
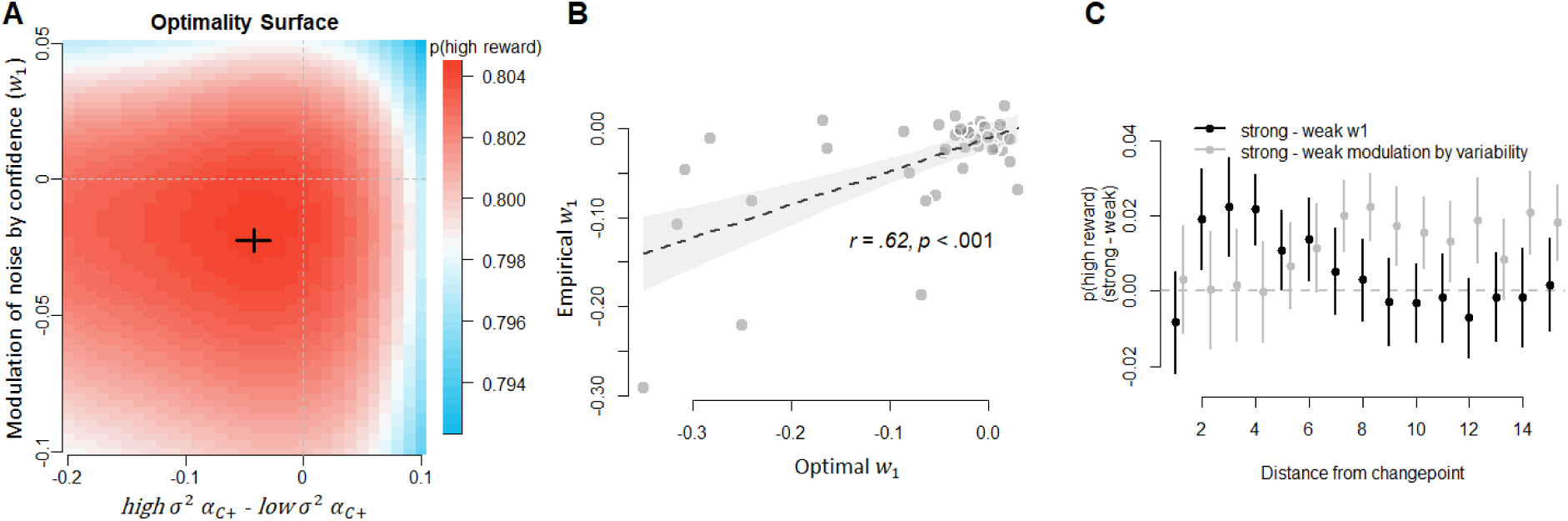
Optimality analysis of the relation between previous-trial confidence and decision noise and outcome variability and α_c+_. **A.** Simulations from Model 16 show that it is optimal to modulate decision noise as a function of previous-trial confidence (i.e. a negative value of w_1_ provides the highest reward) and that it is optimal to have a higher α_c+_ for low vs high outcome variability. Parameter estimates obtained by fitting this model to the empirical data, indicated by the black cross reflecting SEM for both dimensions, lie close to this optimal point. **B.** There was a significant positive relation between the empirically observed w_1_ (controlling the influence of previous-trial confidence on decision noise), and the optimal w_1_. **C.** To explore why it is optimal to modulate decision noise as a function of previous-trial confidence and α_c+_ as a function of outcome variability, empirical data were split into two groups, using a split-median. This analysis revealed the opposite finding for both strategies: Black points shows the difference in learning curves when splitting groups with strong versus weak values of w_1_. The group with a strong modulation of decision noise by previous-trial confidence mostly benefited from this strategy early after a change point but not so much later in time. Grey point show the difference in learning curves when splitting groups according to the modulation of α_c+_ by outcome variability. The group with a strong modulation by outcome variability benefited from this strategy from about trial six onwards, but not early after a changepoint.

First, across participants the value of *w*_1_ that led to the highest p(high reward) was –.015, which would have generated an expected average correct choice of .803. When calculating the optimal *w*_1_ separately for each participant the average optimal point was significantly different from zero, *M* = –.050, *t*(53) = –3.83, *p* < .001. In the data, the average *w*_1_ is –.029, which is very close to the average optimal value (black cross in Figure 5A).

We confirmed that this finding is selective for decision noise, by repeating the same simulations based on Model 10, but now varying the slope controlling for the influence of previous-trial confidence on the learning rate for positive prediction errors. As expected, the optimal value was close to zero, – .0075, and across participants the optimal points did not significantly differ from zero, *p* = .94, *BF* = .15, showing that the link between previous-trial confidence and current trial parameters is selective for decision noise. Thus, we can conclude that it is the optimal strategy to choose the level of decision noise as a function of previous-trial decision confidence, and actual participants closely adhere to this strategy.

Next, we tested in the empirical data whether there was an association between each participant’s optimal value of *w*_1_ and the empirically observed *w*_1_ based on the model fits. As shown in Figure 5B, there was indeed a strong positive correlation between both, *r*(52) = .62, *p* < .001, showing that participants were sensitive to the value of *w*_1_ that maximized their task performance. In line with this finding, participants with a stronger (i.e. more negative) modulation of decision noise by previous-trial confidence also obtain a higher reward, *r*(52) = –.40, *p* = .002. To further unravel the underlying reason for this link, we separately estimated this correlation between modulation and reward based on the first five trials after a changepoint (i.e. a period of high uncertainty) versus the trials from trial six onwards (i.e. a period of low uncertainty). The negative correlation between the modulation of decision noise by previous-trial confidence and reward rate was, again, significant during the period of high uncertainty (i.e. trial 1-5 after a changepoint), *r*(52) = –.60, *p* < .001, but not during a period of low uncertainty (i.e. from trial 6 onwards), *r*(52) = –.20, *p* = .155. This suggests that modulating decision noise as a function of previous-trial confidence is mostly beneficial during periods of high uncertainty (e.g. being noisier after a switch), but not so much during periods of low uncertainty (e.g. excessively sampling the most certain option). To visualize this finding, we computed choice behavior separately for participants with strongly negative *w*_1_ values (strong modulation of decision noise by previous-trial confidence) and participants with a weak *w*_1_ value (weak to no modulation of decision noise by previous-trial confidence), using a median-split analysis. The black dots in Figure 5C shows the difference scores between the learning curves of both groups. As can be seen, participants with a strong modulation of decision noise by confidence more frequently select the high reward option (roughly) between trial 2 and 6 following a switch point, whereas afterwards there is no clear difference between both groups.

Second, we found that it is also optimal to change learning rate *α_c+_* in response to level of outcome variability. When calculating the optimal difference between high and low variability, this difference was significantly different from zero, *M* = –.047, *t*(53) = –3.66, *p* < .001, which is in the range of the empirically observed difference of 0.089 (SD = .24; black cross in 5A). Note that for *α_c+_* there was no correlation between the optimal and the fitted values, *r*(52) = .014, *p* = .920. To unravel why it is optimal to modulate the learning rate as a function of outcome variability, we computed the difference in learning curves between participants with a strong versus weak modulation of *α_c+_* As shown by the grey dots in Figure 5C, the effect is exactly opposite to what was previously reported for *w*_1_: participants with a strong modulation of *α_c+_* more frequently select the high reward option (roughly) starting around 6 trials after a switch point, whereas the first five trials right after the switch there is no clear difference between both groups.

### Transfer effects

The current design dissociated between an induction phase, during which rewards were sampled from a low-variability or a high-variability distribution, and a subsequent test phase, during which rewards were sampled from the same medium-variability distribution. This allowed us to demonstrate that the influence of outcome variability on choices and confidence persists even in the test phase where outcome variability does not differ (i.e. showing a transfer effect). Thus far, however, all models were fitted on the full dataset. To unravel potential transfer effects, we further consider Model 14 (which includes both the effect of outcome variability on the learning rate for positive prediction errors and the effect of previous-trial confidence on decision noise), but we fitted it separately to each phase of the experiment. When fitted to the data of the induction phase only, the results were similar as for the full dataset: there was a significant difference between the learning rate for positive prediction errors in the high versus low variability condition, *M* = .12, *t*(53) = 2.89, *p* = .005, as well as a significantly negative estimate for *w*_1_, *M* = –.033, *t*(53) = –3.72, *p* < .001, showing a modulation of decision noise by previous-trial confidence. Interestingly, when fitting Model 14 to the data of the test phase only, the model still showed the negative estimate for *w*_1_, *M* = –.028, *t*(53) = –3.79, *p* < .001, but the difference between the learning rates was no longer significant, *M* = .05, *t*(53) = 1.70, *p* = .095, *BF* = 0.57. This finding suggests that while outcome variability influenced both choices and confidence during the induction phase, the transfer of these effects on the behavioral level cannot be explained by a transfer in learning rates to the test phase. The implications of these findings as well as a potential explanation are provided in the General Discussion. Finally, the conclusions are identical when estimating Model 6 and Model 13 to both phases of the experiment separately, but these are omitted here for parsimony.

### Causes of confidence in value-based decisions

So far, this paper has treated confidence as the value that participants reported on a scale (i.e. external to the model). However, given that confidence plays a central role in determining decision policy, a natural question is how we can understand confidence within this modeling framework. Given that the current model uses feedback to update its action (*Q*_t_) values on trial *t* associated with each slot machine, a straightforward hypothesis is that confidence is a function of the *Q* values that also inform the choice. To assess this hypothesis, we tested three different models: a model in which confidence only reflects the *Q* value of the chosen option on trial *t* (*Q*_C,t_), a model in which confidence reflects *Q*_C,t_ – *Q*_U,t_, and a model in which confidence reflects 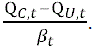. All three models were built on top of the best fitting parameters of Model 14 and included four additional confidence thresholds that mapped the latent value onto a five-point confidence scale (see Methods for details). Given that the three models have the same number of parameters, we directly compared their residual error (see Figure 6A): the data were best explained by a model in which confidence only reflects *Q*_C,t_. The residual error of this model was numerically lower than the differences model, *t*(53) = 1.96, *p* = .055, and the differences scaled by noise model, *t*(53) = 3.18, *p* < .001. This finding is similar to a recent observation made by Salem-Garcia and colleagues (Salem-Garcia et al., 2023), and in line with the finding in perceptual decision making that confidence depends mostly on choice congruent evidence (Maniscalco et al., 2016; Peters et al., 2017; Rollwage et al., 2020; Zylberberg et al., 2012). Model fits of the winning model are shown in Figure 2C-D.

**Figure 6.**
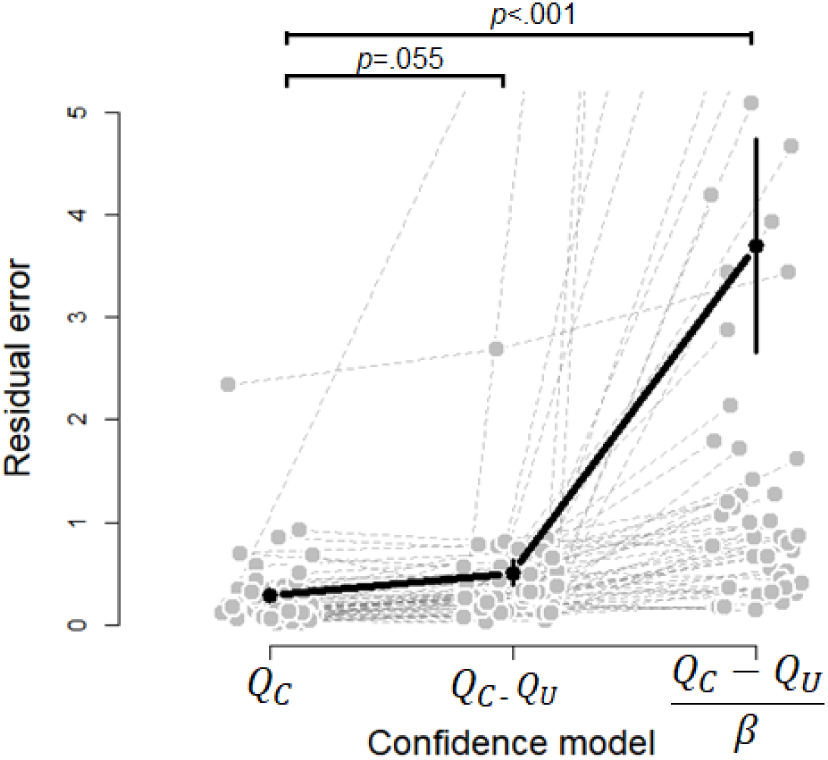
The computational origins of confidence in reinforcement learning. The best model to explain confidence in value-based decisions assumed that confidence was a function of the Q value of the chosen option only. This model outperformed a model assuming that confidence reflects the difference in Q values between the chosen and unchosen option, and a model including decision noise. Fits from this model are shown in Figure 2B-C.

## Discussion

In the current work, we investigated which “meta-signals” guide the process of hyperparameter setting during meta-learning. We attempted to unravel the regulatory role of decision confidence in modulating value learning and decision policy setting, in the context of varying levels of outcome variability. Human participants performed a two-armed bandit task in which they additionally rated the confidence in their decision. Outcome variability was manipulated during an induction phase, and its effect was tested during an unbiased testing phase. Fits from a reinforcement learning model showed a clear dissociation between outcome variability and decision confidence: Outcome variability was involved in value learning, as it influenced the learning rate for positive prediction errors. Previous-trial confidence was involved in setting the decision policy, as it influenced the level of decision noise: when participants reported low versus high confidence in a previous choice they made more noisy (i.e., explorative) versus less noisy (i.e., exploitative) decisions on the current trial. Optimality analysis showed that both strategies are rational approaches which maximize earnings at different temporal loci. The modulation of decision noise by previous-trial confidence was optimal because it induced more frequent exploration early after a changepoint; the modulation of learning rate by outcome variability was advantageous only late after a changepoint because it protected against “chasing the noise” in high variability contexts. Finally, we showed that confidence in two-armed bandit tasks can be modeled as the magnitude of the action value of the chosen option, effectively ignoring the state of the unchosen option. We conclude that variability and confidence have differential roles in meta-learning value learning and decision policy, respectively.

### Meta-signals for meta-learning

Accruing evidence suggests that the hyperparameters controlling learning are not fixed, but instead dynamically vary depending on the environment. This is the case both in optimal computational models (Dayan et al., 2000) as well as in human data (Simoens et al., 2024; Wen et al., 2023). Meta-learning frameworks propose that the setting of hyperparameters governing the learning process is itself a process that is learned (Binz et al., 2023; Doya, 2002; Silvetti et al., 2018). At a fast timescale the brain learns about action values using standard RL algorithms, and the hyperparameters controlling this fast learning are themselves learned at a slow timescale using the same RL algorithms (Silvetti et al., 2018) based on neuromodulatory control (Doya, 2002). More generally, learning at the fast timescale can be formulated as Bayesian inference based on the data, where the precise form of the posterior distribution is learned at the slow timescale (Binz et al., 2023). Meta-learning frameworks make the key prediction that there exist one or more meta-signals that are used by agents to guide this meta-learning process. Similar to how the brain updates its actions value in response to prediction errors (i.e. between expected and obtained rewards), it also updates its hyperparameters based on some meta-signal. Here, we investigated decision confidence, in the context of varying levels of outcome variability, as a prime candidate driving the meta-learning process. Given that (outcome) variability is known to influence subjective confidence (Boldt et al., 2017; Spence et al., 2016) it is unclear whether outcome variability and decision confidence reflect dissociable signals that are differentially used by agents to inform the process of hyperparameter setting. The current work demonstrated a clear dissociation between both, with outcome variability influencing the setting of the learning rate whereas confidence influenced the setting of decision noise. Our findings thus show that humans track and use confidence as a meta-signal that is used to regulate their own learning processes. In the following, we further discuss the role of both decision confidence and outcome variability and their distinctive effects on meta-learning.

### Confidence optimally controls the stochasticity of the decision policy

Participants relied on previous-trial confidence to determine the current level of decision noise. In other words: when participants had low confidence in their decisions they tended to make more explorative choices on the next trial (i.e. exploring whether the other bandit might be more beneficial) whereas when they had high confidence they tended to make more exploitative decisions (i.e. exploiting a known bandit). Our findings are in line with a behavioral study reported by Boldt and colleagues (2019) who showed that participants use confidence in value representations to arbitrate between exploration and exploitation. However, Boldt and colleagues did not estimate computational models, so they were unable to show that confidence interacts with the latent parameter that controls the decision noise. That is exactly what was done in the current work, which showed a close link between decision confidence and decision noise, and we additionally showed that this is in fact the optimal strategy in this task.

From a theoretical perspective, there are (at least) two reasons why this is the optimal strategy; (1) increased exploration around a change point (during which there is high uncertainty), and (2) increased exploitation during stable periods (when there is low uncertainty). First, right after a switch point (subjective) uncertainty is maximal because participants have to figure out whether an unexpected (lack of) reward was caused by a contingency switch or by random noise. In such scenario, it is advantageous to make more exploratory decisions as these will more quickly allow to identify the best bandit. This theoretical scenario is consistent with our empirical data in which we found that participants with a strong link between previous-trial confidence and decision noise mostly benefitted from this during the early period following a rule switch. Note that in our 2-armed bandit task exploration was synonymous to switching to the other bandit, and so future work using larger action spaces should attempt to further arbitrate between directed, undirected and random exploration. Indeed, previous work has shown that uncertainty influences exploration in specific manners (Cogliati-Dezza et al., 2017; Wilson et al., 2014; Wu et al., 2018), so future work should further refine the role of confidence in exploration.

Second, the link between previous-trial confidence and decision noise might be optimal because this allows higher rates of exploitation during periods of low uncertainty. When one is certain that one bandit is the better option, it makes sense to fully exploit that knowledge and always select that bandit. Interestingly, a similar explanation of over-selecting the most valuable bandit was recently used by Lefebvre and colleagues (2022) to explain confirmation bias. Specifically, the observation that humans have larger learning rates for positive than for negative prediction errors (for chosen options; i.e., a confirmation bias), maximizes reward rate relative to an unbiased updating rule. Although both explanations are very similar at the outcome level (i.e. both low decision noise and higher learning rates for positive prediction errors lead to a subtle over-selection of the most valuable bandit), at the computational level the implementation is radically different. In the case of previous confidence affecting the level of decision noise, action values themselves are not affected. In contrast, in the case of a confirmation bias action values about the bandits are affected (i.e. because of the difference in learning rate for positive and negative prediction errors), but the level of decision noise that is used to translate these into a choice is not affected.

Finally, although we did not find compelling evidence in the current work that confidence influences the learning rate, this does not imply that confidence plays no role in learning (for theoretical discussions, see Cortese, 2022; Drugowitsch et al., 2019; Meyniel et al., 2015). Previous work has shown that in the context of perceptual learning, confidence prediction errors shape the learning process (Balsdon et al., 2023; Guggenmos et al., 2016), and similar to “classical” prediction errors such confidence prediction errors have been observed in the striatum (Daniel & Pollmann, 2012). A notable difference with the current work is that in these studies participants did not receive external feedback, thus forcing them to rely on internal feedback signals to regulate learning (see also Frömer et al., 2021). Finally, in the context of a probability learning task, it has been shown that confidence serves as a weighting factor that balances prior beliefs and environmental feedback (Meyniel et al., 2016; Meyniel & Dehaene, 2016), suggesting a link between confidence and the learning rate.

### Outcome variability optimally affects learning, unmediated by confidence

Given that outcome variability is known to influence the learning rate (Diederen & Schultz, 2015; Preuschoff & Bossaerts, 2007), and variability is known to influence confidence (Boldt et al., 2017; Spence et al., 2016), one potential hypothesis was that confidence mediates the influence of outcome variability on the learning rate. Although we replicated both these findings, we did not find compelling evidence that confidence modulates the learning rate. Consistent with this, results from a mediation analyses showed that although outcome variability affects confidence, its effect on choice behavior was not mediated by confidence. Thus, although outcome variability does influence confidence, both variables have distinct effects on hyperparameter setting. The observation that outcome variability influences the learning rate for positive prediction errors, is in line with previous work showing that increased uncertainty leads to reduced learning rates, which is the optimal strategy under this scenario (Behrens et al., 2007; Preuschoff & Bossaerts, 2007). This strategy was optimal because during “stable” periods (i.e. late after a changepoint) as it prevented overshooting action values in response to prediction errors that are caused by noise at the outcome level.

### Transfer effects of outcome variability

In order to empirically establish meta-learning, a common approach is to induce a certain hyperparameter setting in an induction phase and then test whether the manipulation transfers to an unbiased testing phase. For example, Wen and colleagues (Wen et al., 2023) showed that after performing a probabilistic card sorting task in a high or low volatility context, this strategy transferred into an unbiased testing phase during which volatility was matched (see also Simoens et al., 2024). We used a highly similar approach in which rewards were drawn from high or low variability distributions during an induction phase, and during a subsequent test phase rewards were drawn from the same medium-variability distribution. At the behavioral level there was clear evidence for a transfer effect: both during the induction and the test phase the learning curves were steeper when rewards were sampled from the low variability distribution; a finding that was mirrored in confidence judgments. Intriguingly, however, the model fits did not support a straightforward interpretation in terms of transfer of hyperparameters. Although there was a clear modulation of the learning rate for positive prediction errors by the level of outcome variability during the induction phase, this effect was not significant during the test phase. The influence of previous-trial confidence on decision noise, however, was significant in both induction and test phase. One potential explanation for this finding is that outcome variability influences confidence (as shown in the empirical data), which then affected the learning curves in test phase only indirectly via its influence on decision noise. Finally, we note that although the current work describes key evidence for the possibility of confidence reflecting a meta-signal that guides the process of meta-learning, we did not actually implement meta-learning into our computational framework. Future work should attempt to implement yet another meta-level where the optimal value of w_1_ is learned by the model (i.e. rather than estimated, as done in the current work), and likewise for the optimal modulation of learning rate by outcome variability.

### Data and code availability

All experiment code, analysis code and raw data have been deposited online and can be freely accessed [insert link upon publication].

## Acknowledgments

The authors like to thank Monica De Bock for help with data collection and Senne Braem, Sara Ershadmanesh and Alexandre Lietard for useful comments on an earlier draft. This research was supported by a Horizon Europe MSCA Doctoral Networks (COnfident DEcisions; CODE).

